# A Novel Propidium Monoazide-Based PCR Assay Can Measure Viable Uropathogenic *E. coli in Vitro* and *in Vivo*

**DOI:** 10.1101/2021.10.13.464198

**Authors:** Albert S Lee, Olivia K Lamanna, Kenji Ishida, Elaise Hill, Michael H. Hsieh

## Abstract

**Background:** Polymerase chain reaction (PCR) is an important means by which to study the urine microbiome and is emerging as possible alternative to urine cultures to identify pathogens that cause urinary tract infection (UTI). However, PCR is limited by its inability to differentiate DNA originating from viable, metabolically active versus non-viable, inactive bacteria. This drawback has led to concerns that urobiome studies and PCR-based diagnosis of UTI are confounded by the presence of relic DNA from non-viable bacteria in urine. Propidium monoazide (PMA) dye can penetrate cells with compromised cell membranes and covalently bind to DNA, rendering it inaccessible to amplification by PCR. Although PMA has been shown to differentiate between non-viable and viable bacteria in various settings, its effectiveness in urine has not been previously studied. We sought to investigate the ability of PMA to differentiate between viable and non-viable bacteria in urine.

**Methods:** Varying amounts of viable or non-viable uropathogenic *E. coli* (UTI89) or buffer control were titrated with mouse urine. The samples were centrifuged to collect urine sediment or not centrifuged. Urine samples were incubated with PMA and DNA cross-linked using blue LED light. DNA was isolated and uidA gene-specific PCR was performed. For *in vivo* studies, mice were inoculated with UTI89, followed by ciprofloxacin treatment or no treatment. After the completion of ciprofloxacin treatment, an aliquot of urine was plated on non-selective LB agar and another aliquot was treated with PMA and subjected to uidA-specific PCR.

**Results:** PMA’s efficiency in excluding DNA signal from non-viable bacteria was significantly higher in bacterial samples in phosphate-buffered saline (PBS, dC_T_=13.69) versus bacterial samples in unspun urine (dC_T_=1.58). This discrepancy was diminished by spinning down urine-based bacterial samples to collect sediment and resuspending it in PBS prior to PMA treatment. In 3 of 5 replicate groups of UTI89-infected mice, no bacteria grew in culture; however, there was PCR amplification of *E. coli* after PMA treatment in 2 of those 3 groups.

**Conclusion:** We have successfully developed PMA-based PCR methods for amplifying DNA from live bacteria in urine. Our results suggest that non-PMA bound DNA from live bacteria can be present in urine, even after antibiotic treatment.

This indicates that viable but non-culturable *E. coli* can be present following treatment of UTI, and may explain why some patients have persistent symptoms but negative urine cultures following UTI treatment.

## INTRODUCTION

The existence of the urinary microbiome, the presence of bacterial communities within the urinary tract, is challenging the paradigm that this organ system is normally sterile (Siddiqui et al., 2012; Wolfe et al., 2012; Hilt et al., 2014; Brubaker and Wolfe, 2015; Thomas-White et al., 2016). Furthermore, several studies have shown an association between the urine microbiome and numerous urological diseases (Fouts et al., 2012; Siddiqui et al., 2012; Whiteside et al., 2015; Bajic et al., 2018; Bučević Popović et al., 2018; Magistro and Stief, 2019; Neugent et al., 2020). Therefore, it is imperative to accurately characterize the urinary microbiome as it may inform overall urinary tract health and aid in the diagnosis of urinary conditions, i.e., urinary tract infection (UTI) (Perez-Carrasco et al., 2021).

Numerous clinical studies of patients with UTI feature assessments of both microbiologic and clinical cure, which are based on negative urine cultures and resolution/improvement of symptoms, respectively (Raz et al., 2002; Wunderink et al., 2018; Miller et al., 2019). Some patients in these studies have featured discordance between microbiologic and clinical cure (Hilt et al., 2014; Price et al., 2018; Swamy et al., 2019). One possible interpretation of this discordance is that conventional urine cultures may be missing residual bacteria causing persistent symptoms following antibiotic therapy. An alternative to urine cultures for detection of urinary microorganisms is polymerase chain reaction (PCR). PCR identifies organisms through the amplification of DNA material present in urine and many studies on the urinary microbiome rely on this molecular approach (Lewis et al., 2013; Brubaker and Wolfe, 2015; Ackerman et al., 2019). These methods do not discriminate between relic DNA (DNA from non-viable bacteria) versus DNA from viable bacteria. This is an important limitation of conventional PCR because the confounding effects of relic DNA have been reported in various microbiologic settings (Carini et al., 2016; Nagler et al., 2021; Ren et al., 2021). Viable, metabolically active bacteria presumably exert much more influence over the clinical course of UTI than dead bacteria. Thus, the amplification of total DNA without selection for DNA from viable bacteria may bias conventional PCR-derived results (Carini et al., 2016; Lennon et al., 2018). Given that relic DNA influences conventional molecular measurements of microbial abundance and diversity, we posit that a method to detect viable, metabolically active bacteria is needed for more accurate urobiome studies (Lennon et al., 2018).

A method of identifying metabolically active bacteria via PCR has been recently developed. Propidium monoazide (PMA) dye penetrates cells with compromised cell membranes (non-viable cells) and covalently binds to DNA, rendering it unable to be amplified by PCR (Deshmukh et al., 2020). PMA-based PCR has previously been shown to differentiate between non-viable and viable bacteria in many settings (Fittipaldi et al., 2010; Cattani et al., 2016; Gobert et al., 2018; Brauge et al., 2019; Lu et al., 2019). However, the efficiency of PMA in urine has not been previously investigated. Here we investigated the ability of PMA dye in urine to detect DNA derived from viable bacteria.

## MATERIALS AND METHODS

### Bacteria Culture

All bacteria work was performed under sterile conditions in a BSL-2 biosafety cabinet. Bacteria was prepared by previously reported methods (Hung, 2009). Briefly, glycerol stock containing the uropathogenic *Escherichia coli* strain UTI89 was used to inoculate a Miller Luria Broth (LB) agar plate (Sigma-Aldrich, St. Louis, MO). The plate was incubated for 24 hours at 37°C. A single colony was picked and transferred to 10 mL LB broth. The culture was incubated overnight at 37°C in a stationary flask. Twenty-five microliters of the 10 mL culture were transferred to 25mL of LB broth. The culture was incubated again overnight at 37°C in a stationary flask. The culture was centrifuged at 5000 x g for 5 minutes at 4°C. The supernatant was decanted and the bacteria pellet was resuspended in 10 mL of sterile phosphate buffered saline (PBS) (Thermofisher Scientific, Waltham, MA). This suspension was diluted tenfold in sterile PBS. The optical density (OD) 600nm value was analyzed using the NanoDrop-1000 (Thermofisher Scientific, Waltham, MA) and the suspension diluted until the OD was 0.50, corresponding to 1-2×10^7^ colony-forming units (CFU) per 50 μL.

### Mouse Urine Collection

Mice were scruffed and held with their pelvises above sterile parafilm (Sigma-Aldrich, St. Louis, MO) until they voided. Urine was aspirated from the parafilm. New parafilm was used for each mouse. Urine was then placed on ice and immediately processed.

### Generation of non-viable bacteria

Five hundred μL of *E. coli* with OD value 0.5 was mixed with isopropanol (Sigma-Aldrich, St. Louis, MO) to achieve a final concentration of 70% v/v. After 10 minutes, the mixture was centrifuged at 8000 x g for 10 min. The supernatant was removed and the pellet was resuspended in 100 μL of PBS. The suspensions of non-viable bacteria were plated on LB with agar plates and incubated at 37°C overnight to confirm successful killing.

### Urine dilution of bacteria

Mouse urine was serially diluted to a ratio of 1:2, 1:4, 1:8, and 1:12 with PBS. Subsequently, 50 μL of all viable or all non-viable bacteria was added to 50 μL of the various titrations of urine, or undiluted urine. The samples were then either treated with PMA or left untreated.

### PMA treatment

Under minimal light, PMAxx Dye (hereafter referred to as PMA) (Biotium, Fremont, CA) with a concentration of 20 mM was diluted with nuclease-free water (Sigma-Aldrich, St. Louis, MO) to a final concentration of 10 mM. It was then added to the bacterial mixture in a 1:100 ratio. Next, samples were incubated for 15 minutes in the dark with gentle agitation. The samples were placed in an LED lightbox (Biotium, Fremont, CA) for 20 minutes to induce PMA crosslinking of DNA. The supernatant was removed and the pellet was reconstituted with PBS to its original volume of 100 μL.

### DNA extraction and quantification

DNA was isolated using the DNeasy PowerSoil Pro Kit (Qiagen, Germantown, MD) according to kit instructions, except that DNA was eluted from the column with 25 μL of nuclease-free water. PCR was performed targeting the *E. coli uidA* gene with TaqMan polymerase (Invitrogen, Waltham, MA) according to previously described methods (Taskin et al., 2011). Delta C_T_ (dC_T_) values were calculated to quantify the differences between C_T_ values of PMA-treated and untreated samples.

### Preparation of urea solution and PMA treatment

Urea (Sigma-Aldrich, St. Louis, MO) was diluted with PBS in to two concentrations; 285 mM, corresponding to the urine urea level in humans and 1800 mM, corresponding to the urine urea level in mice (Yang and Bankir, 2005). Fifty μL of 100% viable or 100% non-viable bacteria was added to 50 μL of the urea solutions. The DNA of the PMA treated and untreated samples was extracted and amplified as described above.

### Titration of viable and non-viable bacteria with urine to develop a standard curve

Defined quantities of viable bacteria, isopropanol-killed non-viable bacteria, or PBS were mixed to a total volume of 100 μL. With a fixed amount of viable bacteria, non-viable bacteria were added to achieve 1:10, 1:100 and 1:1000 non-viable to viable bacterial dilutions. In a similar manner, various amounts of viable bacteria were added to a fixed amount of non-viable bacteria to generate a standard curve. Undiluted viable and non-viable cultures were also used. Fifty μL of these bacterial solutions were added to urine. The mixture was then centrifuged at 5000xg, resuspended with 100 μL of sterile PBS, treated with PMA, and the DNA was extracted as outlined above.

### Bacterial inoculation via transurethral catheterization of mice

All animal work was approved by The Institutional Animal Care and Use Committee of Children’s National Hospital under Animal Use Protocol #00030764. Procedures were performed in an ethical fashion. Prior to use, all animals were acclimated for 7 days after arrival to the animal facility. 24-week-old female C3H/HeOuJ mice (stock no: 000635, The Jackson Laboratory, Bar Harbor, ME) were used in this study.

Mice were anesthetized using 2% isoflurane. Any urine in the bladder was expressed by gently pressing on the lower abdomen. A 24g x 3/4 inch angiocatheter (Clint Pharmaceuticals, Old Hickory, TN) was attached to a prepared 1 ml syringe containing the inoculant. The angiocath was lubricated (DynaLub Sterile Lubricating Jelly, Amazon, Seattle, WA) and transurethrally inserted into the bladder. 100 μL of the inoculant was instilled slowly into the bladder and the angiocatheter kept inserted for 30 seconds to prevent leakage of the inoculant.

### Antibiotic treatment of mice

Five days after inoculation, mice were intraperitoneally injected with 10 mg/kg ciprofloxacin twice a day. This regimen was selected as it recapitulates the human plasma peak levels achieved with the commonly used 500 mg oral dose, and has been shown previously to adequately treat UTI in mice (Guillard et al., 2013).

### Urine collection for in vivo studies

One day after completion of ciprofloxacin treatment, mouse urine was collected on ice. Individual urine samples in the same treatment groups (3-4 mice/group) were pooled. The urine was either serially diluted and plated in triplicate on LB agar or prepared for PMA treatment. Fifty μL of PBS was added to the urine and the solution was centrifuged and treated with PMA as outlined above. DNA was extracted and the E. coli *uidA* gene was amplified as described above (Taskin et al., 2011).

## RESULTS

### Urine interferes with PMA efficiency

Initial PMA-based PCR experiments using mouse urine yielded little differences in amplification of PMA-treated vs. untreated DNA samples. This led us to consider the possibility that urine was exerting a matrix effect which interferes with downstream molecular processes such as PMA crosslinking (Taylor et al., 2012). The dC_T_ value of PMA-treated vs. untreated samples that contained 100% non-viable bacteria resuspended in mouse urine was 1.58, which was about one tenth of the dC_T_ of the same sample resuspended in PBS (13.69, **Figure 1A**). When urine was diluted with PBS, the dC_T_s of PMA-treated vs. untreated samples increased, indicating improved PMA efficiency (**Figure 1B**). However, the increase in dCT plateaued at a dilution of 1:8. These findings indicate that urine inhibits PMA activity.

**Figure 1 (A-D):**
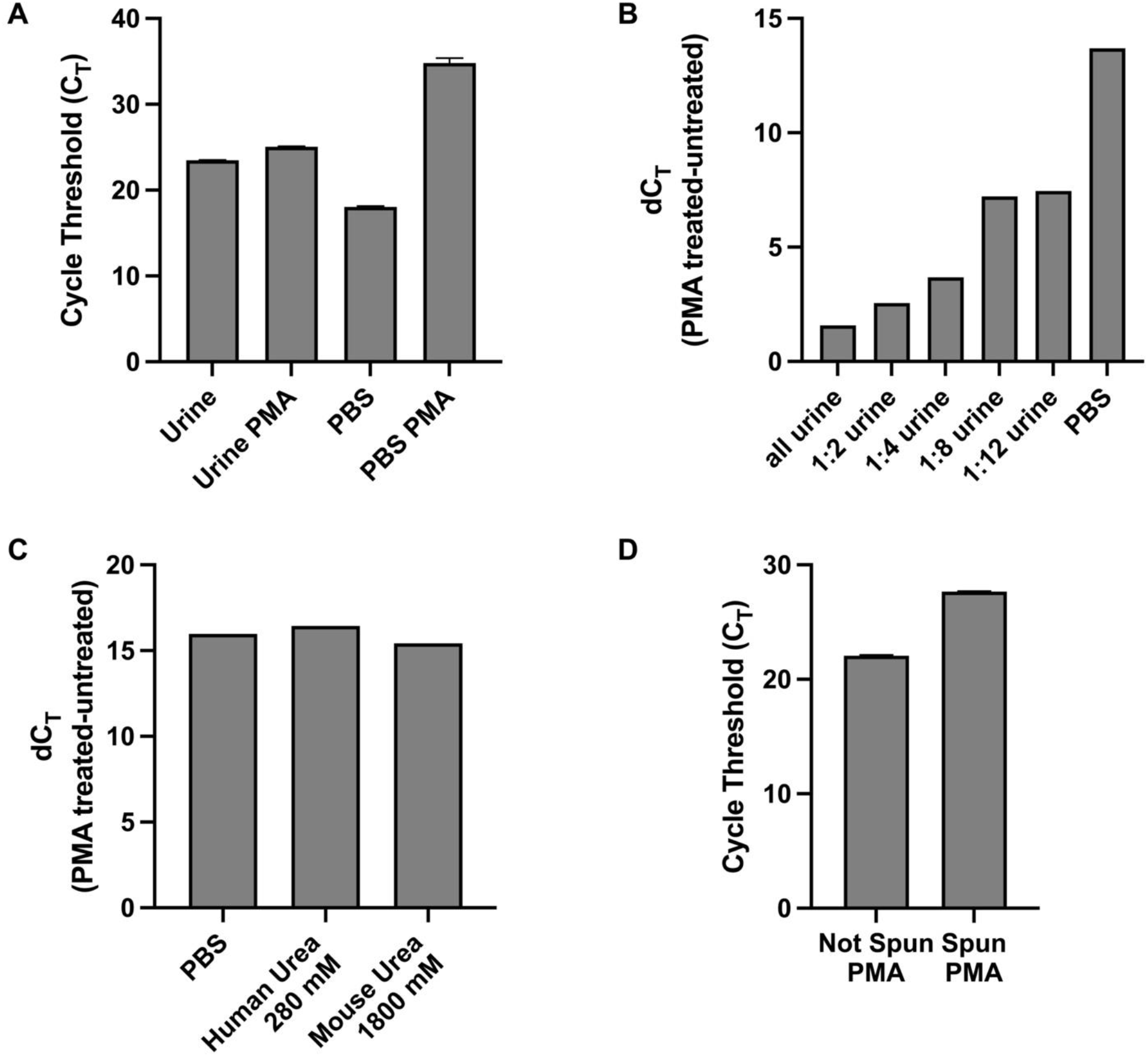
The matrix effect of urine on PMA crosslinking of DNA is not urea-based and is eliminated by centrifugation and resuspension in PBS. A) C_T_ values of PMA-treated and untreated samples of non-viable bacteria resuspended in mouse urine or PBS. B) The difference in C_T_ (dC_T_) values of nonviable bacteria resuspended in various dilutions of urine or PBS. C) C_T_ values of non-viable bacteria treated with PMA in PBS or urea concentrations corresponding to human and mouse urine. D) C_T_ values of non-viable bacteria in urine with or without resuspending the contents in PBS before PMA treatment. Data shown is representative of three replicate experiments.

### Urea does not affect PMA efficacy

Urea is the most abundant solute present in urine and is known to influence molecular structure and function (Yang and Bankir, 2005; Wei et al., 2010). Thus, we sought to investigate urea’s potential effect on PMA’s function in crosslinking DNA and subsequently inhibiting its amplification. We analyzed PMA’s efficiency at two different urea concentrations: 280 mM and 1800 mM, the approximate concentration of urea in human and mouse urine, respectively. The dCTs of PMA-treated vs. untreated samples with 100% nonviable bacteria suspended in either urea concentration was similar to that of the PBS control. Namely, the dCT of the PBS, human urea concentration, and mouse urea concentration samples were 15.98, 16.42, and 15.43, respectively (**Figure 1C**). This suggests that urea does not cause urine’s matrix effect on PMA efficiency.

### Resuspension of urine sediment with PBS restores PMA efficiency

We observed that for a solution with 100% nonviable bacteria, the C_T_ value increases when the urine supernatant is resuspended in PBS prior to PMA crosslinking compared to when PMA crosslinking is performed in unspun urine (**Figure 1D)**. Furthermore, upon removal of the urine supernatant and subsequent resuspension in PBS, PMA treatment and downstream PCR was most efficient in differentiating viability when there was a greater proportion of nonviable cells in the solution. Across titrations of viable and non-viable bacteria where viable bacteria make up the majority of the solution, C_T_ values did not significantly differ, with all values close to 16. This suggests the amount of non-viable bacteria does not influence the detection of a fixed amount of viable bacteria. Conversely, when the majority of the cells are non-viable, the C_T_ values decrease as the amount of viable bacteria in the solution increases. For instance, the C_T_ is approximately 18 with 100% viable cells in solution and increases to ∼ 27 when viable cells make up only 0.1% of the solution (**Figure 2B**).

**Figure 2 (A,B):**
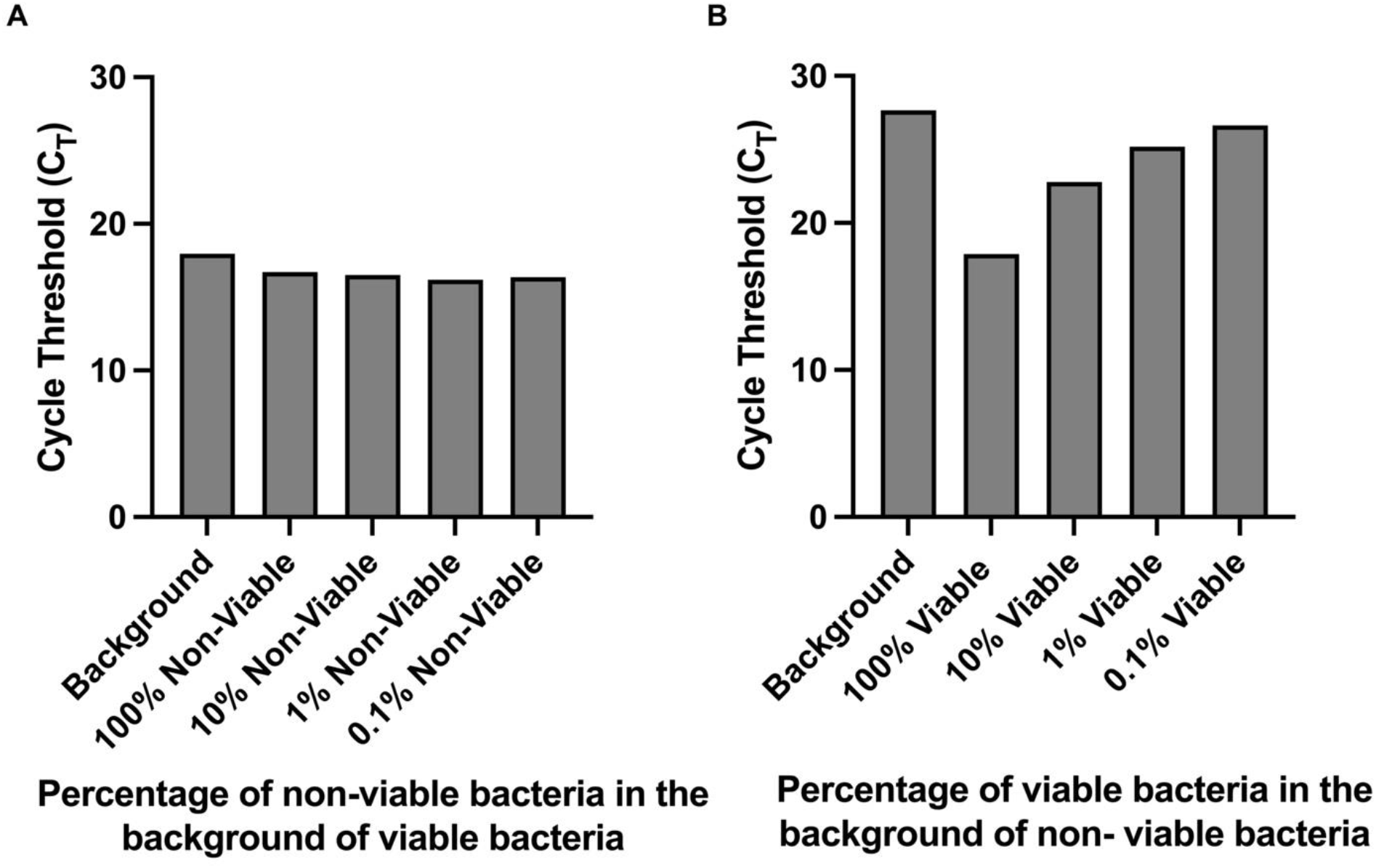
PMA crosslinking of DNA is most efficient when most of the bacteria is non-viable. Bacterial samples were centrifuged and resuspended in PBS before PMA treatment. A) C_T_ values of samples with a fixed amount of viable bacteria and titrated amounts of nonviable bacteria. B) C_T_ values of samples with a fixed amount of non-viable bacteria and titrated amounts of viable bacteria.

### Detection of viable bacteria correlates with colony forming units (cfu)

Given that the number of colony forming units present in urine cultures remains the mainstay of clinical diagnosis of UTI, we sought to determine whether urine cultures with various titrations of viable cells and a fixed amount of nonviable cells yielded cfu and C_T_ values that correlated with each other. Indeed, the correlation between cfu and C_T_ values was strongly negative with an *r*^2^ of 0.955 (**Figure 3**).

**Figure 3:**
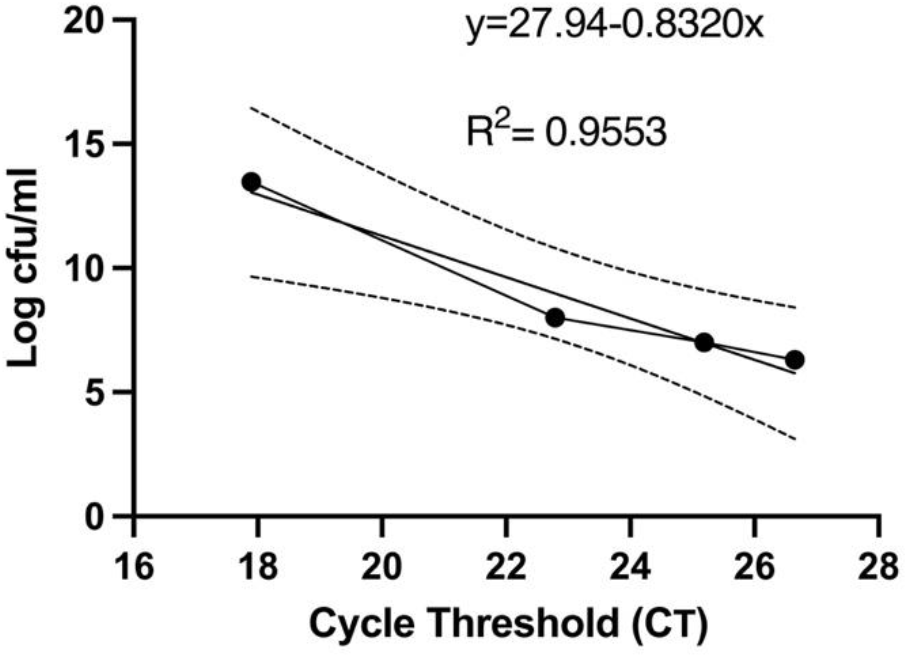
Colony forming units and PMA-based viability PCR cycle threshold number has a strong negative correlation. Before PMA treatment, an aliquot of bacterial samples was serial diluted, plated on LB agar plates, and cfu were counted after 24 hours. cfu per ml were plotted against the respective C_T_ values. The linear regression and 95% confidence interval band is shown. All experiments were repeated 3 times.

**Figure 4:**
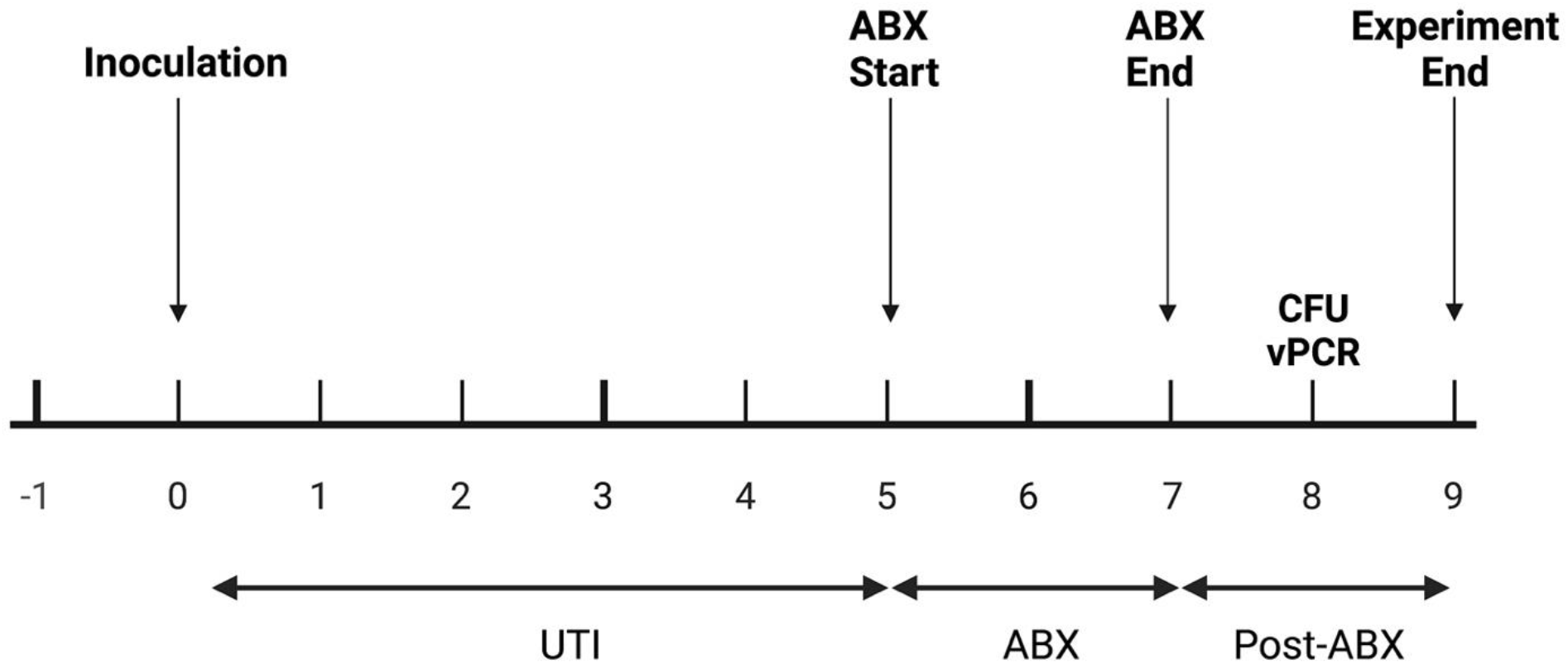
In vivo experimental plan. Mice were inoculated with UTI89 via transurethral catheterization on day 0 and given ciprofloxacin (“ABX”) twice daily on days 5, 6, and 7 post-infection. On day 8, one day after the completion of antibiotics, urine was collected to measure cfu/mL and to perform PMA-based viability PCR (“vPCR”).

### Detection of non-culturable but viable bacteria in mouse urine after antibiotic treatment

PCR-based detection of bacterial DNA in urine from patients with persistent UTI symptoms and negative cultures following antibiotic therapy has been criticized as being confounded by the presence of relic DNA (Lehmann et al., 2011). To investigate whether non-culturable, viable bacteria can still be present in urine after antibiotic treatment of UTI, we administered uropathogenic *E. coli* (UTI89) to mice and treated them with ciprofloxacin according to established protocols (Hung, 2009). One day after the completion of antibiotic treatment, 3 out of the 5 replicate groups had no bacterial growth on non-selective LB agar (**Figure 5**). However, after PMA treatment of these samples, PCR successfully amplified the *E. coli uidA* gene in 2 out of the 3 culture-negative groups, indicating the presence of viable, nonculturable bacteria. Based on our standard curve (**Figure 3**), these 2 groups contained 1 and 6×10^5^ cfu/ml *E. coli*.

**Figure 5:**
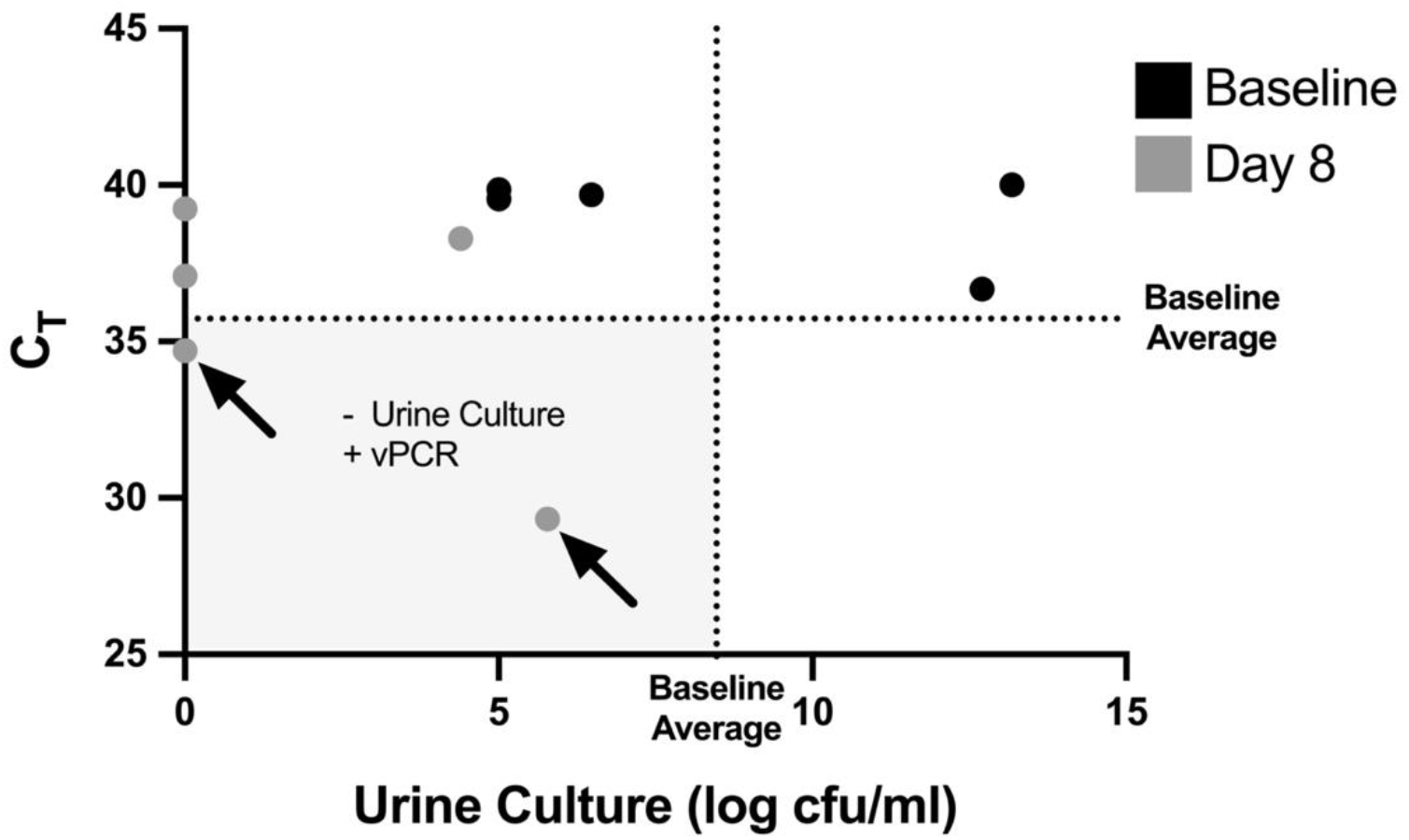
Non-culturable, live bacteria detected in mouse urine by PMA-based PCR after antibiotic treatment. Mice were given UTIs on day 0 and were administered ciprofloxacin starting on day 5 and ending after day 7 (see Figure 4). Graph depicts C_T_ values and log transformed urine cfu/mL values at baseline before inoculation (black dots), and at day 8 after antibiotic treatment (grey dots). Each dot represents a pooled cohort of 2-3 mice. The average CT and log cfu/ml of the baseline samples are indicated by the dotted lines. The greyed quadrant (Q3) represents values that are considered as having a negative urine culture and positive PMA-based viability PCR (“vPCR”). Data shown is pooled from two set of experiments.

## DISCUSSION

Our findings indicate that a PMA-based urine PCR is an appropriate method to distinguish viable and non-viable *E. coli* in the urine for both *in vitro* and *in vivo* applications. We were able to eliminate the signal from soluble relic DNA and DNA from nonviable cells. By resuspending urine contents in PBS before PMA treatment we established an easy and reproducible method to eliminate soluble relic DNA while preserving *E. coli* cells. This approach yielded dC_T_ values similar to that of non-urine exposed *E. coli* resuspended in PBS and those reported in the literature (Taskin et al., 2011). Thus, we demonstrate a novel method to utilize PMA in urine that will allow for PCR-based studies to selectively identify viable bacteria in urine.

Our preliminary experiments pointed to a matrix effect of murine urine that inhibited PMA crosslinking of DNA (Chamberlain et al., 2019). We initially focused on urea as a potential cause of this effect. Urea is a by-product of amino acid metabolism and one of the most abundant urine solutes. Mouse urine has a higher urea concentration compared to human urine, which led us to speculate that urea could be influencing PMA crosslinking function in mouse urine (Yang and Bankir, 2005). The similar dC_T_ of *E. coli* in urea vs. PBS suggested urea is not the substance in urine that inhibits PMA’s crosslinking efficiency. Thus, the inhibitory effect of urine is likely is due to a non-urea-related matrix effect. The simple steps of centrifuging bacteria-containing samples and resuspending them in buffer may eliminate any matrix effect of other biofluids and environmental samples of interest, enabling PMA-based PCR amplification of DNA from viable bacteria in other settings.

Identifying viable, but potentially unculturable bacteria may improve understanding of bacterial biology in patients with recurrent UTI. We identified non-culturable but viable *E. coli* in the urine of infected mice given ciprofloxacin. Non-culturable but viable bacteria in settings other than the urinary tract are a recognized phenomenon (Coutard et al., 2005; Oliver, 2005). However, the presence of such bacteria in urine is poorly characterized. It may be that these bacteria represent intracellular *E. coli* which have formed intracellular communities and become quiescent reservoirs of infection (Mulvey et al., 2001; Rosen et al., 2007). Our findings may explain why, despite patients having undergone susceptibility-guided antibiotic treatment and a subsequent negative test-of-cure by urine culture, some of these patients experience recurrent UTI.

Compared to urine culture, PMA-based urine PCR has the clinical advantages of a more rapid time to UTI diagnosis and broader organism detection. While enhanced quantitative urine culture has demonstrated greater sensitivity for uropathogen detection than conventional culture (Price et al., 2016), it is still time-intensive. In contrast, a uropathogen-specific PCR platform based on PMA could detect multiple viable organisms quickly.

This is a preliminary study using a new molecular method for identification of bacteria in urine. A potential limitation to our study is the use of PMA dye prior to PCR. Specifically, studies have shown that PMA results can be skewed by specific primers. However, the primers used in this study have been shown to be effective in multiple studies for *E. coli* without loss of viability data (Taskin et al., 2011; van Frankenhuyzen et al., 2013). Finally, it remains to be determined how this PMA-based PCR platform performs with human urine.

## Abbreviations

CFU: Colony forming units
C_T_: Threshold Cycle
dC_T_: Delta Threshold Cycle
*E. coli*: *Escherichia coli*
LB: Luria-Bertani
OD: Optical Density
PBS: Phosphate-buffered saline
PMA: Propidium Monoazide
RT-PCR: Real-time polymerase chain reaction
UTI: Urinary Tract Infection

## ACKNOWLEDGMENTS

This work was supported by NIH-R01DK113504 (MH) and George Washington University Cancer Biology Training Program NIH-T32CA247756 (KI)

## CONTRIBUTION TO THE FIELD STATEMENT

Conventional PCR detection of bacteria in the urine cannot differentiate between viable and non-viable bacteria. Here, we describe a novel PCR-based method to selectively detect DNA from viable bacteria in urine. In mice given urinary tract infections, we demonstrate that viable bacteria still remain in urine after a complete course of antibiotics. This finding may account for recurrent UTI or persistent symptoms in some patients treated with antibiotics for UTI who have negative post-treatment urine cultures. Compared to conventional urine cultures, this PCR-based method may be superior in sensitivity for the diagnosis of UTI.

## Notes

### Competing Interest Statement

The authors have declared no competing interest.

